# Chromosome-level genome assembly of the European leaf-toed gecko, *Euleptes europaea*

**DOI:** 10.64898/2026.08.10.744031

**Authors:** Josephine R. Paris, Linelle Abueg, Sarah Pelan, Ying Sims, Tatiana Tilley, Jacquelyn Mountcastle, Jennifer Balacco, Brian O’Toole, Olivier Fedrigo, Giulio Formenti, Erich D. Jarvis, Daniele Canestrelli, Daniele Salvi

## Abstract

The European leaf-toed gecko (*Euleptes europaea*) is a small, nocturnal gecko endemic to the western Mediterranean. As a phylogenetically distinctive member of the Gondwanan family Sphaerodactylidae, it represents an important species for studying Mediterranean island biogeography, adaptation, and reptile genome evolution. The species also occupies a key position for investigating the evolution of sex chromosomes, as geckos exhibit remarkable diversity and frequent transitions in sex-determination systems. We present a chromosome-level genome assembly of *Euleptes europaea* generated as part of the Vertebrate Genomes Project. The 1.8 Gb assembly has a scaffold N50 of 102.3 Mb (contig N50 27 Mb), with 21 chromosome-scale scaffolds corresponding to the known karyotype (2n = 42). The primary assembly has a BUSCO completeness of 97.80% (95.60% as single-copy), a *k*-mer completeness of 96.00%, and a *k*-mer quality value (QV) of 61.20. Repetitive elements account for 53.20% of the genome and genome annotation identified 18,633 protein-coding genes. This high-quality reference genome will facilitate studies of genome evolution, island adaptation, and sex chromosome evolution across geckos and other reptiles.

## Background & Summary

*Euleptes europaea* (the European leaf-toed gecko or in Italian *tarantolino*) is a small (SVL: 30-40 mm; ^1,2^), strictly nocturnal gecko endemic to the western Mediterranean. It is the sole representative of the monotypic genus *Euleptes* ^*3*^ and belongs to the Gondwanan family Sphaerodactylidae, which is otherwise distributed primarily throughout the Neotropics ^4^. This makes *E. europaea* a remarkable biogeographical relict within the European fauna. The species occurs across Sardinia and Corsica together with nearly all of their satellite islands; the mainland and islands of southern France; the coast and two islets of Liguria in north-western Italy; the coast of Tuscany and the Tuscan Archipelago; and three islands off the coast of Tunisia ^5^.

Well adapted to insular environments, *E. europaea* is frequently the only resident vertebrate on small offshore islets, where it occupies rocky crevices in Mediterranean habitats. Although considered a rock-specialist, recent reports have documented arboreal behaviour across a wide variety of trees, shrubs, and bushes ^6^. As one of the smallest European geckos, the species is characterised by cryptic colouration, adhesive toe and tail pads, and limited dispersal ability. Owing to its fragmented insular distribution, phylogenetic distinctiveness, and ecological specialisation, *E. europaea* is emerging as a valuable model for investigating island biogeography, population divergence, and adaptation in Mediterranean reptiles ^1,6–9^. Despite its ecological and evolutionary interest, genomic resources for the species have remained limited.

Geckos are characterised by exceptional lability in sex determination, with repeated transitions between environmental and genotypic systems, as well as among XX/XY and ZZ/ZW chromosomal systems across the clade ^10–12^. However, many lineages remain poorly characterised, particularly within the ancient family Sphaerodactylidae, where both male and female heterogametic systems have been reported ^11,13^. Cytogenetic analyses of *E. europaea* have suggested a possible XX/XY system based on subtle male chromosomal heteromorphism ^14^, but the genomic basis of sex determination in the species remains unresolved. As the sole representative of an early-diverging genus, and the species with the highest diploid chromosome number currently reported among sphaerodactylids, *E. europaea* occupies a key phylogenetic position for understanding the tempo and mode of sex chromosome evolution in geckos.

Here, we present a chromosome-level genome assembly of the European leaf-toed gecko, *Euleptes europaea* (Fig. 1), assembled as part of the Vertebrate Genomes Project (VGP) ^15^ using PacBio HiFi sequencing, Bionano optical maps and Arima Hi-C technology. The assembly spans 1.8 Gb, with a scaffold N50 of 102.3 (contig N50: 27 Mb), arranged across 21 chromosome-scale scaffolds (Fig. 2; Table 1), consistent with the known karyotype of the species (2n = 42) ^14^. Macrosynteny was near one-to-one with *Teratoscincus roborowskii* (family Sphaerodactylidae), with the higher chromosome number of *E. europaea* (21 versus 18) being attributable to three lineage-specific fissions, which are also supported by comparison with a more distant outgroup *Heteronotia binoei* (family Gekkonidae) (Fig. 3). The primary assembly (rEulEur1.hap1) has a BUSCO score of 97.80%, a *k*-mer completeness of 96.00%, and a *k*-mer QV value of 61.18. Repetitive elements comprise 53.20% of the genome. We predicted 18,633 protein-coding genes, with a BUSCO completeness score of 96.60% and an OMArk completeness score of 98.32%. Beyond providing a high-quality genomic resource for a phylogenetically distinctive Mediterranean reptile, this assembly establishes a framework for future studies aimed at identifying sex-determining regions, reconstructing the history of sex chromosome turnover, and investigating genome evolution in geckos.

**Table 1.** Genome assembly statistics.

| Quality metric | rEulEur1.hap1 | rEulEur1.hap2 |
| --- | --- | --- |
| Estimated genome size (bp) | 1,589,842,322 | 1,589,842,322 |
| Size of final assembly (bp) | 1,781,916,107 | 1,775,520,415 |
| Number of chromosomes | 21 | 0 |
| Number of scaffolds | 570 | 381 |
| Contig N50 (Mb) | 27 | 32 |
| Contig L50 | 20 | 19 |
| Scaffold N50 (Mb) | 102.3 | NA |
| Scaffold L50 | 7 | NA |
| Average scaffold length (bp) | 2,419,987 | NA |
| Largest scaffold (bp) | 98,702,465 | NA |
| GC content (%) | 44.68 | 44.50 |
| Repeat content (%) | 53.20 | NA |
| *k*-mer base pair QV | 61.18 | 61.73 |
| *k*-mer completeness (%) | 96.04 | 95.90 |
| BUSCO | C:97.8%[S:95.6%,D:2.2%],F:0.2%,M:2.1%,n:6118,E:8.4% | C:97.6%[S:95.3%,D:2.4%],F:0.2%,M:2.2%,n:6118,E:8.3% |

**Figure 1.**
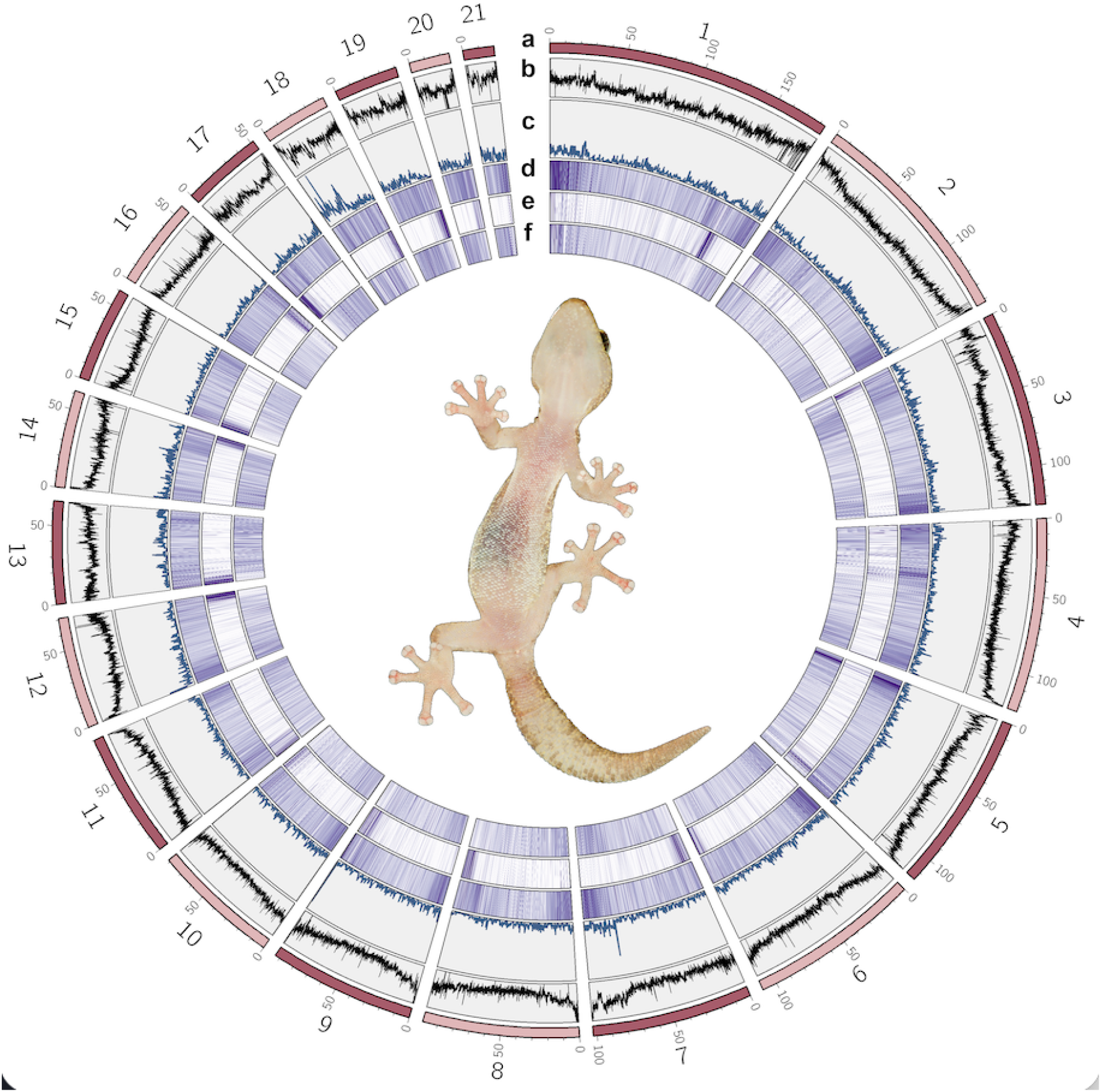
Genomic landscape of the *Euleptes europaea* chromosome-level reference genome (rEulEur1; GCF_029931775.1). Circos plot summarising the distribution of genomic features across the 21 assembled chromosomes (∼1.75 Gb of the 1.78 Gb assembly). Tracks, from outermost to innermost, show: (**a**) the chromosome ideograms with a physical scale in megabases (Mb); (**b**) GC content in 50-kb windows (black line); (**c**) gene density, expressed as the number of genes per 200-kb window (blue line), based on the NCBI RefSeq annotation (20,622 protein-coding and non-coding genes on chromosomes); and the density of the three most abundant transposable-element classes in 50-kb windows, shown as heatmaps: (**d**) LINEs (∼18.4% of the genome), (**e**) LTR retrotransposons (∼5.1%), and (**f**) SINEs (∼4.8%). Repeats were annotated with EarlGrey; interspersed repeats occupy ∼53% of the genome overall. In the heatmap tracks, colour intensity scales from low (white) to high (dark purple) repeat density; each track is scaled independently to its own range to aid visualisation, so intensities are not directly comparable between tracks (d–f). Centre: an adult *E. europaea*. Photograph used with permission from Bram Conings ©.

**Figure 2.**
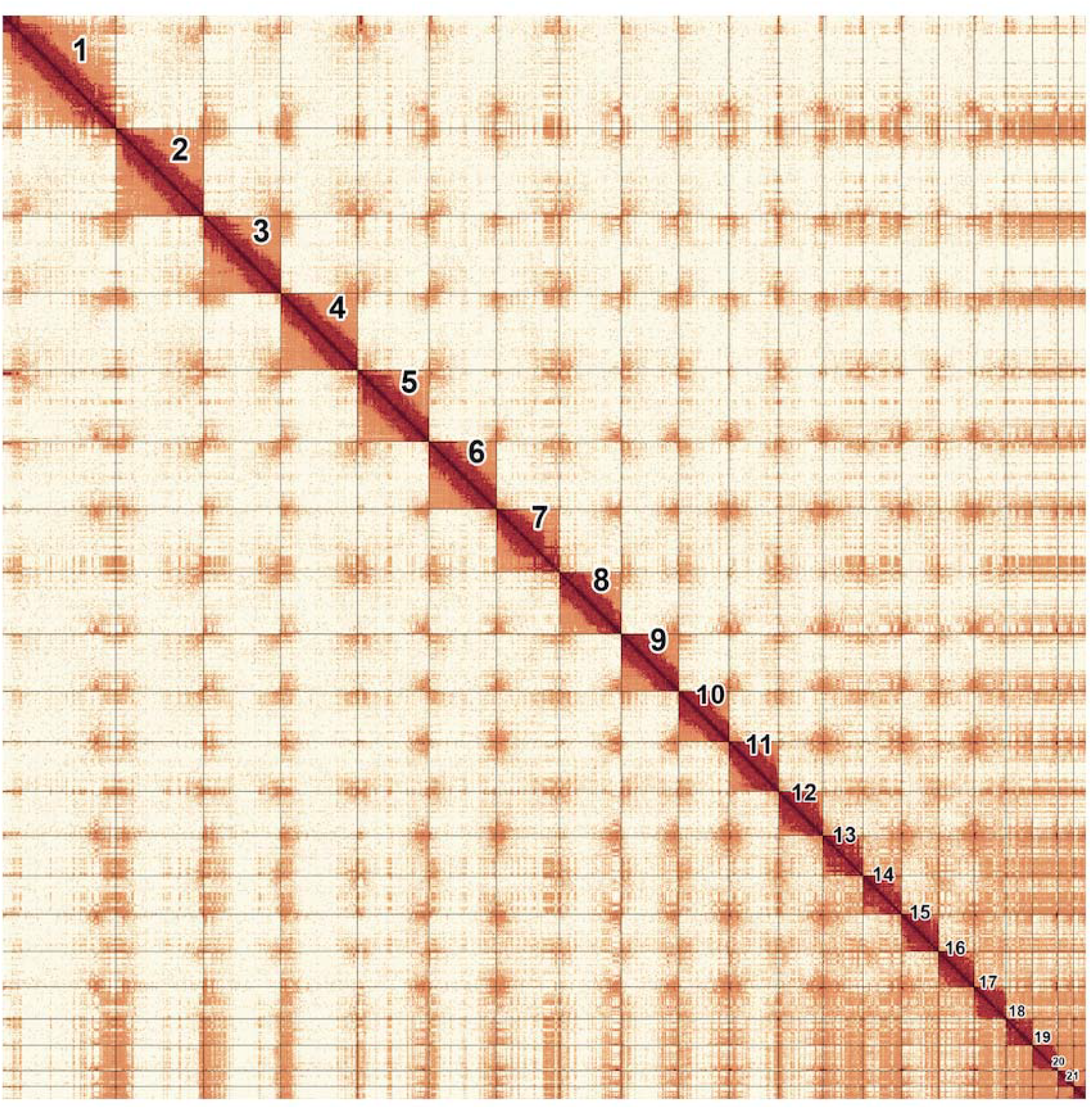
Hi-C contact map of the curated *Euleptes europaea* assembly (rEulEur1). Genome-wide Hi-C contact matrix after manual curation, showing the 21 chromosome-scale scaffolds (numbered by decreasing size along the diagonal). Contact frequency is coloured from low (pale) to high (dark red); the strong on-diagonal signal within each block and the sharp boundaries confirm correct chromosome scaffolding. Image generated with PretextView/PretextSnapshot.

**Figure 3.**
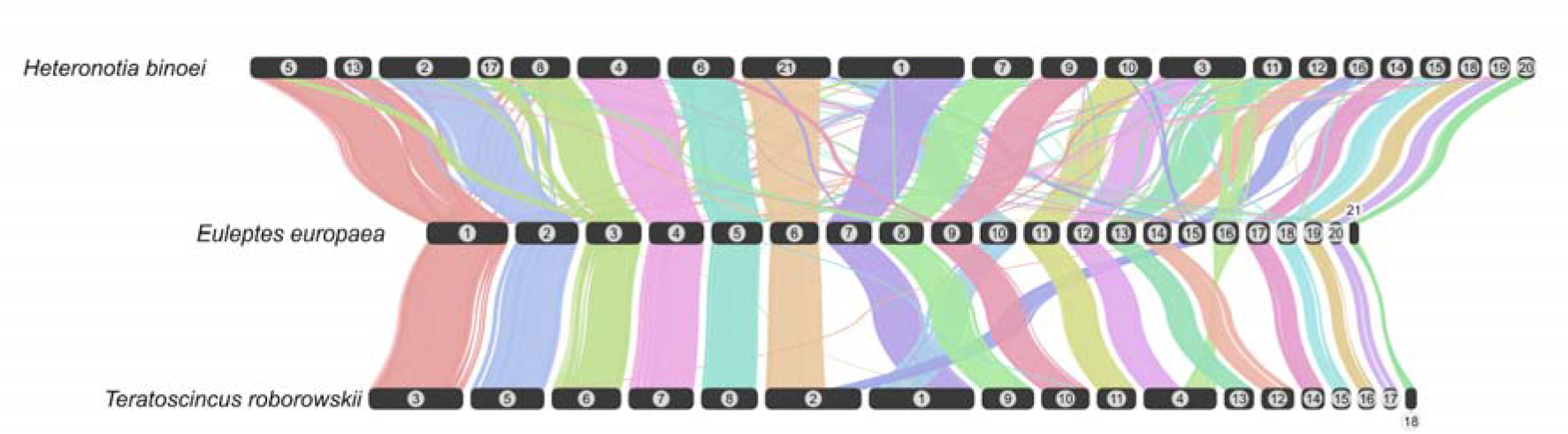
Macrosynteny between *Euleptes europaea* and two other gekkotan genomes. Chromosomes are drawn to scale as dark bars and labelled with their assembly chromosome number: *Heteronotia binoei* (top; GCF_032191835.1, 21 chromosomes), *Euleptes europaea* (middle; rEulEur1.hap1, GCF_029931775.1, 21 chromosomes) and *Teratoscincus roborowskii* (bottom; GCA_051473905.1, 18 chromosomes). Ribbons connect homologous blocks and are coloured by the *E. europaea* chromosome they involve. In the two flanking assemblies, chromosomes have been reordered to follow the *E. europaea* order. Crossing ribbons indicate inversions or rearrangements relative to *E. europaea* rather than differences in assembly conventions.

## Methods

### Sample collection, extraction, and sequencing

The genome sample was obtained from an adult male collected in mainland Corsica (42.249549 N 8.660302 E) under the authorisation approval of the Prefét de la Corse-du-Sud (n° 2A-2019-04-18-005). The specimen was anaesthetised prior to dissection. Skin and intestinal tissues were excluded, and the remaining tissues were flash-frozen and stored at -80°C.

For PacBio, HMW DNA was isolated from skeletal muscle using the MagAttract HMW DNA Kit (Qiagen 67563). A total of 38 mg of frozen tissue was disrupted with a Qiagen TissueRuptor II (Cat. No. 9002755). After the tissue homogenization, lysis and subsequent DNA isolation was performed following the protocol described in the MagAttract HMW DNA Handbook (Manual Purification of High-Molecular-Weight Genomic DNA from Fresh or Frozen Tissue). The purified DNA was eluted in 100□µL of Qiagen Buffer AE. The DNA was quantified with triplicate measures using a Qubit 3 fluorometer (Invitrogen Qubit dsDNA Broad Range Assay cat no. Q32850). Prior to HiFi library preparation, the DNA was sheared using the Megaruptor 3 (Diagenode, Denville, NJ, USA). HiFi libraries were prepared with sheared DNA and the SMRTbell Express Template Prep Kit 2.0 (Pacific Biosciences, Menlo Park, CA, USA). Size-selection was performed with the Pippin HT (Sage Science, Beverly, MA, USA) with a 10kb size cut-off. The libraries were sequenced on a PacBio Sequel IIe, with Sequencing Plate 2.0 and 8M SMRT cells, and Binding Kit 2.2, generating a total of 21 Gb of data.

For the Bionano data, HMW DNA was extracted from skeletal muscle with the Circulomics Nanobind Tissue Big DNA Kit. The DNA was quantified using the Qubit 3 fluorometer (Invitrogen Qubit dsDNA Broad Range Assay cat no. Q32850) and fragment size was assessed with a pulsed field gel electrophoresis (Pippin Pulse, SAGE Science, Beverly, MA). 750 ng DNA was labeled using direct labeling enzyme (DLE1) and the Bionano Prep Direct Label and Stain (DLS) protocol (document number 30206) and then imaged on the Bionano Saphyr instrument, generating 880 MiB of data (∼190X coverage).

For the Hi-C libraries, 37 mg of skeletal muscle was used for the Arima Genomics crosslinking reaction following the manufacturer’s low input sample amount guidance (Arima High Coverage HiC Kit Document Part Number: A160162). Libraries were prepared using the Arima-HiC 2.0 kit (Arima Genomics, CA, USA). The library was sequenced with the Illumina NovaSeq 6000 platform with 150□bp paired-end reads, generating a total of 252 Gbp of data (∼99X coverage).

### Genome assembly

The genome was assembled using the Vertebrate Genomes Project (VGP) v2.1 Galaxy pipeline ^16^. Genome characteristics were estimated from *k*-mer profiles generated with Meryl ^17^ , and analysed with GenomeScope v2 ^18^. HiFi sequences and Hi-C data were used as input to assemble phased contigs using HiFiasm v0.16.1 in Hi-C mode ^19^. Haplotypes were scaffolded using the Bionano and Hi-C contact data. Bionano scaffolding was achieved using Bionano Solve v3.7.0 ^20^ with default parameters and without contig breaking. Hi-C scaffolding was performed on the Bionano scaffolds. Hi-C reads were aligned and prepared for scaffolding using the Arima mapping pipeline, which uses bwa mem ^21^ and samtools ^22^ for mapping and filtering. Scaffolding was performed using YaHS v1.2 ^23^. PretextMap (https://github.com/wtsi-hpag/PretextMap) was used to visualise Hi-C contacts before and after scaffolding. Haplotypes were manually curated with gEVAL ^24^ using PretextMaps to correct potential assembly structural errors, to manually join and align unplaced scaffolds ^25^.

### Transposable element annotation

Repetitive elements were identified and annotated with EarlGrey v7.2.6 ^26^ which integrates *de novo* repeat discovery, automated curation, and homology-based annotation. A species-specific repeat library was generated *de novo* with RepeatModeler v2.0.7 ^27^ with NCBI/RMBLAST 2.10.0+ against the Dfam v3.9 ^28^ Vertebrata partition. Consensus sequences were refined through EarlGrey’s iterative “BLAST, Extract, Extend” curation process; redundant families were clustered with CD-HIT v4.8.1 ^29^. The curated library was combined with known repeats and used to soft-mask the assembly with RepeatMasker v4.1.7 ^30^. TE libraries were clustered to reduce redundancy (-c) and spurious TE annotations□<□100□bp were removed (-m). Overlapping and nested annotations were resolved to produce non-redundant repeat coordinates. Repetitive sequences span 947.96 Mb, or 53.2% of the assembly. Transposable elements account for the majority of this content, dominated by LINEs (18.4%), with smaller contributions from LTR retrotransposons (5.1%), SINEs (4.8%), DNA transposons (1.5%), and Penelope-like elements (1.5%). A further 18.9% of the genome comprised repeats that could not be assigned to a known class, and 3.9 % consisted of simple repeats, satellites, and other low-complexity sequences.

### Gene prediction and functional annotation

Gene prediction and functional annotation was performed by the National Center for Biotechnology Information (NCBI) using the NCBI Eukaryotic Genome Annotation Pipeline v10.1 (EGAP; ^31^) on the rEulEur1.hap1 assembly ^32^. BUSCO v6.0.0 analysis was performed in protein mode to assess quality, using the sauropsida_odb12 (n□=□6,118) OrthoDb v12 dataset ^33^. OMArk v0.3.0 ^34^ using OMAmer v2.0.3 was run using the Lepidosauria clade (12,331 HOGs), using the longest isoform of each protein. In total, 21,512 genes and pseudogenes were predicted, including 18,633 protein-coding genes (18,276 fully-supported mRNAs), and 2,325 non-coding RNAs.

### Mitogenome assembly and annotation

The mitogenome was assembled using MitoHifi v2 ^35^, using MitoFinder v1.4.2 ^36^ for annotation and mitoVGP 2.2 ^37^ for quality control. The mitogenome of Przewalski’s wonder gecko (*Teratoscincus przewalskii*; NC_067620) was used as the starting sequence. The resulting mitogenome was 17,541 bp in length and contained the standard 37 vertebrate mitochondrial genes (13 protein-coding, 22 tRNAs, and 2 rRNAs).

### Genome synteny analysis

To explore genome synteny (Fig. 3), we compared the genome of *E. europaea* against two other chromosome-level gecko assemblies: the closer-related *Teratoscincus roborowskii* (Sphaerodactylidae; GCA_051473905.1) and a more distant outgroup, *Heteronotia binoei* (Gekkonidae; GCF_032191835.1). Macrosynteny was visualised with NGenomeSyn v1.43 ^38^.

Synteny with *T. roborowskii*, which lacks a genome annotation, was inferred at the nucleotide level by aligning the two genomes with minimap2 v2.17 ^39^ using the asm20 preset for divergent genomes. Primary alignments (tag tp:A:P) ≥ 2 kb with mapping quality ≥ 5 were retained and chained into syntenic blocks: collinear alignments sharing a chromosome pair and strand and separated by ≤ 500 kb on both genomes were merged, and merged blocks ≥ 150 kb were kept (293 blocks spanning 1.63 Gb of the *E. europaea* assembly).

Synteny with the more divergent *H. binoei* was inferred from protein-coding gene order using the RefSeq protein and annotation (GFF) sets for *E. europaea* (GCF_029931775.1) and *H. binoei* (GCF_032191835.1). For each gene we kept the longest protein isoform and restricted the set to chromosomally placed genes (*E. europaea*, 18,461; *H. binoei*, 20,224). All-versus-all homology was computed using DIAMOND v2.0.5 blastp ^40^ (e-value ≤ 1 × 10^− 10^, --max-target-seqs 5; 152,460 hits), and collinear blocks were identified with MCScanX ^41^ with default parameters. The 233 inter-specific collinear blocks were converted to syntenic links, with block orientation determined from the sign of the covariance between paired gene midpoints in the two genomes.

Macrosynteny between *E. europaea* and *T. roborowskii* was essentially one-to-one: each of the 21 *E. europaea* chromosomes had a single *T. roborowskii* counterpart accounting for ≥95% of its syntenic sequence, and all 18 *T. roborowskii* chromosomes were represented. The difference in chromosome number was accounted for by three *T. roborowskii* chromosomes that each corresponded to two *E. europaea* chromosomes (Tro1 = Eeu7 + Eeu10; Tro2 = Eeu6 + Eeu15; Tro4 = Eeu12 + Eeu16). Synteny with the more distant *H. binoei* was correspondingly more fragmented, but these same pairs were largely co-located in this assembly (most clearly Eeu7 and Eeu10, which placed 94.7% and 92.9% of their syntenic sequence on *H. binoei* chromosome 1).

### Data Records

Raw sequencing and mapping data are available from the VGP GenomeArk repository (https://www.genomeark.org/genomeark-curated-assembly/Euleptes_europaea.html) and on the NCBI/ENA under BioProject: PRJNA919166 (https://www.ncbi.nlm.nih.gov/bioproject/919166).

The primary genome assembly (rEulEur1.hap1) is available at NCBI GenBank under the accession GCF_029931775.1 ^32^. It is also available on the UCSC Genome Browser (https://genome.ucsc.edu/h/GCF_029931775)

The alternate haplotype (rEulEur1.hap2) is available at NCBI GenBank under the accession GCA_029931755.1 ^42^ It is also available in the UCSC Genome Browser (https://genome.ucsc.edu/h/GCA_029931755.1).

The mitochondrial genome sequence is available in NCBI GenBank, accession JARABC010000592 ^43^

### Technical Validation

Genome profiling using *k*-mers (*k* = 21) provided an estimated genome size of 1,589,842,322 bp with a heterozygosity of 0.92%. The rEulEur1.hap1 assembly is 1,781,916,107 bp in length and the rEulEur1.hap2 assembly is 1,775,520,415 bp in length. Assembly quality checks were performed using Merqury v1.3 ^17^, gfastats ^44^, and BUSCO v6.0.0 with the sauropsida_odb12 (n□=□6,118) OrthoDb v12 dataset ^33^. The BUSCO score of the rEulEur1.hap1 assembly is 97.8% complete (95.6% as single-copy, 2.2% as duplicated), 0.2% fragmented, 2.1% missing. The BUSCO score of the rEulEur1.hap2 assembly is 97.6% complete (95.3% as single-copy, 2.4% as duplicated), 0.2% fragmented, 2.2% missing. The Merqury *k*-mer assessment revealed a QV score of 61.18 for rEulEur1.hap1 and 61.73 for rEulEur1.hap2 (QV = 61.44 for both haplotypes). Merqury completeness was 96.04% for rEulEur1.hap1 and 95.90% for rEulEur1.hap2 (99.43% for both haplotypes). We found that the majority (98.35%) of the assembled genome is contained within the 21 largest scaffolded chromosomes confirmed by Hi-C analysis.

To assess annotation quality and completeness, we used BUSCO (same dataset as above for the genome) in protein mode and OMArk v0.3.0 ^34^ using the Lepidosauria clade (12,331 HOGs). BUSCO completeness of the proteome is: 96.6% complete (94.3% as single-copy, 2.3% as duplicated), 0.8% fragmented, 2.6% missing. OMArk completeness is 98.32%, including 96.41% single and 1.92% duplicated (of which 13% were expected and 1.80% were unexpected). There were 1.68% missing HOGs.

## Code Availability

All software and pipelines were executed according to the methods section. No custom code was generated for this study.

## Author Contributions

D.S. conceived the study; D.S. collected the sample; D.S. and D.C. contributed anatomical sampling of tissues and exported the isolated samples for sequencing and genome assembly at the Vertebrates Genome Laboratory, The Rockefeller University; T.T. performed DNA isolation, J.M. generated Bionano optical maps, B.O. generated the Hi-C data, J.B. generated the PacBio data with supervision from O.F.; L.A. assembled the genome with supervision from G.F. and E.D.J.; Y.S. generated the curation data sets; S.P. performed manual curation of assembled primary sequences; J.R.P. assessed the assembly and annotation quality, performed repeat annotation, performed synteny analyses, created the figures and wrote the manuscript. All authors read, edited, and approved the final manuscript.

